# Increased functional coupling of the mu opioid receptor in the anterior insula of depressed individuals

**DOI:** 10.1101/2020.10.04.325449

**Authors:** Pierre-Eric Lutz, Daniel Almeida, Dominique Filliol, Fabrice Jollant, Brigitte L. Kieffer, Gustavo Turecki

## Abstract

The mu opioid receptor (MOR) is a G protein-coupled receptor that plays an essential role in reward and hedonic processes, and that has been implicated in disorders such as depression and addiction. Over the last decade, several brain imaging studies in depressed patients have consistently found that dysregulation of MOR function occurs in particular in the anterior insular cortex, an important brain site for the perception of internal states and emotional regulation. To investigate molecular mechanisms that may underlie these effects, here we assessed genetic polymorphisms, expression, and functional G-protein coupling of MOR in a large post-mortem cohort (N=95) composed of depressed individuals who died by suicide, and healthy controls. Results indicated that depression, but not comorbid substance use disorder or acute opiate consumption, was associated with increased MOR activity. This effect was partly explained by a specific increase in expression of the inhibitory alpha G-protein subunit GNAI2. Consistent with previous neuroimaging studies, our findings support the notion that enhanced endogenous opioidergic tone in the anterior insula may buffer negative affective states in depressed individuals, a mechanism that could potentially contribute to the antidepressant efficacy of emerging opioid-based medications.

## Introduction

Depression is a recurring and often chronic brain disorder that stems from maladaptive processes affecting multiple neurotransmitter systems. While the monoaminergic theory of depression received considerable attention, and fuelled intense research efforts over the last 50 years, another theory, dating back to the early 1900s, posits the implication of the opioid system in regulating mood[1]. Euphoric properties of opiates acting at the mu opioid receptor (MOR), such as morphine and heroin, have been recreationally exploited for centuries, and were once considered a potential ‘opioid cure’ for depression.

Over the last two decades, both clinical and animal research have further strengthened the notion that adaptations at the level of MOR signalling may contribute to depression pathophysiology [1]. Recently, functional neuroimaging has allowed studies of MOR in patients experiencing various mood states [2]. Positron emission tomography (PET) with the MOR selective radiotracer [^11^C]-carfentanil found that the induction of a sadness state, or of sustained pain, was associated in both healthy [3,4] and depressed [5] women with changes in MOR bioavailability across several brain regions. Among these regions, the insular cortex [6] appears as a major site where MOR modulates affective states, in particular in relation to social experiences, such as social acceptance and rejection, in both healthy volunteers [7] and depressed [8] subjects.

While PET imaging allows studying MOR availability in living individuals, this approach does not differentiate changes in occupancy of the receptor by endogenous opioid peptides from changes in receptor abundance, complicating interpretation of findings. Since MOR belongs to the large family of inhibitory G protein-coupled receptors (GPCR), radioligand binding assays in post-mortem cohorts offer a complementary approach to assess receptor level and signalling. The five previous studies available for MOR in relation to depression were conducted between 1990 to 2012 [9–13], focused on relatively small cohorts, and investigated the temporal and frontal cortices, caudate nucleus, and thalamus. Importantly, no investigation was conducted on the cortical structure most consistently identified by the aforementioned brain imaging studies, the insular cortex.

In this context, the goal of the present work was to test the hypothesis that major depressive disorder (MDD) may associate with dysregulated MOR function in the anterior insular cortex (AI). Using post-mortem samples from a large and well-characterized cohort (N=95 total), we compared subjects diagnosed with MDD (N=55), and healthy controls (N=40). Expression of MOR was measured using quantitative PCR, while MOR coupling was assessed using agonist-stimulated [^35^S]-GTPγS binding, and the expression of Gi proteins measured using Nanostring. In addition, a series of single nucleotide polymorphisms located along the MOR gene and potentially contributing to variability in receptor expression or function were characterized. Overall, our results indicate that MDD associates in the anterior insula with increased MOR coupling, which appears partly mediated by increased expression of the alpha(i2) G-protein subunit, suggesting a molecular substrate that may contribute to disease pathophysiology.

## Material and methods

### Cohort

This study was approved by the Research Ethics Board at the Douglas Mental Health University Institute, Verdun, Quebec. Brain tissue was obtained from the Douglas-Bell Canada Brain Bank (http://douglasbrainbank.ca/). We compared two groups of subjects matched for sex, age, post-mortem interval (PMI) and pH (see Table 1): 1) One group of individuals who died by suicide during a major depressive episode (MDD group), and 2) A second group of psychiatrically healthy individuals who died suddenly from accidental causes, and who constituted the control group (C). Information on psychopathology, socio-demographic variables and development was obtained by way of psychological autopsies performed by trained clinicians, with the informants best acquainted with the deceased, as described elsewhere [14]. The process employed by our group in the collection of information used in psychological autopsies has been extensively investigated and found to produce valid information, especially in the context of observable behaviours and major life events [15–17]. Both cases and controls were characterized by the same psychological autopsy methods, avoiding the occurrence of systematic biases. Psychopathology diagnoses were obtained using DSM-IV criteria by means of SCID-I interviews adapted for psychological autopsies. Brain samples were dissected from each subject from 0.5 cm-thick coronal brain sections by expert brain bank staff following standard dissection procedures, with the aid of a human brain atlas (see [18] and thehumanbrain.info/brain/bn_brain_atlas/brain.html). Samples were carefully dissected at 4°C after having been flash-frozen in isopentene at −80°C. AI tissue was dissected from the superior part of the insula, just beneath the overlying opercula in the brain section containing the striatum, at the level of nucleus accumbens.

**Table 1.** Sample Demographics. Healthy controls (C) and subjects who died by suicide during a major depressive episode (MDD) were matched for sex, age, post-mortem interval (PMI), and pH. There were no differences, between groups, in the number of male and female subjects, the presence of alcohol or drug abuse, or RIN values (see main text). Age, RIN, PMI and pH values are shown as mean ± sem.

|  | Controls (C) | Major depressive disorder (MDD) |
| --- | --- | --- |
| N | 40 | 55 |
| Sex, M/F (%) | 32 / 8<br>(80.0 / 20.0%) | 45 / 10<br>(81.8 / 18.2%) |
| Age (years) | 45.65 ± 2.98 | 45.73 ± 2.15 |
| RNA integrity number (RIN) | 6.89 ± 0.16 | 7.15 ± 0.14 |
| PMI (hours) | 27.3 ± 3.52 | 25.8 ± 2.51 |
| pH | 6.44 ± 0.05 | 6.56 ± 0.04 |
| Alcohol/Drug Abuse | 14<br>(35.0%) | 20<br>(36.4%) |

### Relationship to previous studies

Our previous study on a partly overlapping but smaller cohort (N=59) focused on the kappa opioid receptor [19] and its epigenetic regulation, while also reporting mRNA expression for the MOR. The present study investigated an enlarged cohort (N=95), and comprehensively characterized MOR for gene expression, genetic variation, and signalling efficiency.

### Quantitative polymerase chain reaction (qPCR)

RNA was extracted from homogenized brain tissue samples using the RNeasy Lipid Tissue Mini Kit (Qiagen). RNA quantity and quality were measured by Nanodrop® and Agilent 2100 Bioanalyzer technologies, and only samples with an RNA integrity number (RIN) ≥ 5 were used (resulting in N=40 C and N=55 MDD subjects). Extracted RNA was reverse-transcribed using M-MLV reverse transcriptase (Invitrogen™). Messenger RNA (mRNA) levels were quantified by real time polymerase chain reaction (RT-PCR) using SYBR® Green DNA intercalating dye and master mix (Bio-Rad) and the ABI 7900HT PCR machine. Primers for targeted genes were designed using Primer-BLAST (http://www.ncbi.nlm.nih.gov/tools/primer-blast/) and validated by gel migration and dissociation curves. Complementary DNAs (cDNA) from every subject were pooled and used to prepare calibration curves, from which cDNA quantities of target genes were calculated in each sample according to the measured RT-PCR threshold cycle (Ct value), in quintuplicates. Relative expression levels for each gene of interest were calculated by dividing the cDNA quantity for the gene of interest by the arithmetic mean of cDNA quantities for two reference housekeeping genes, GAPDH and β-actin. MOR, forward primer: TTC CGT ACT CCC CGA AAT GC; reverse primer: ACA ATC TAT GGA ACC TTG CCT G. GAPDH, forward primer: TTG TCA AGC TCA TTT CCT GG; reverse primer: TGT GAG GAG GGG AGA TTC AG. β-actin, forward primer: AAG ACC TGT ACG CCA ACA CA; reverse primer: GCA GTG ATC TCC TTC TGC ATC.

### Genotyping

Genomic DNA extracted from brain tissue was first amplified, via PCR, using a Platinum Taq DNA Polymerase (Invitrogen®). Forward and reverse primers were designed using NCBI’s Primer-BLAST tool and synthesized by Integrated DNA Technologies (IDT). Final reactions contained 1 μL of gDNA, 1 μL of 10× PCR buffer, 0.2μL of 10mM dNTP mix, 0.6μL of 50mM MgCl2, 0.1μL of Q-solution, 1μL of 10nM forward and reverse primers, 0.1μL of DNA Polymerase, and 6μL of H2O. PCR reactions were carried out using the following conditions: 95°C for 5 min, then 35 cycles of [95°C for 30s, 60°C for 30s, 72°C for 40s], final extension at 72°C for 7 min. Individual amplicons were then purified using a 0.7x AMPure bead mix. Purified amplicons were sent for Sanger sequencing, on a 3730xl DNA Analyzer (Applied Biosystems), at the McGill University and Genome Quebec Innovation Center. Sequence trace files in ABI format, as well as reference sequences in FASTA format, were loaded into the novoSNP 3.0.1 program for Windows users (Weckx et al., 2005). Variation at each location was determined by following the procedures outlined in Weckx et al [20].

### Agonist-stimulated [^35^S]-GTPγS binding activity of MOR

The functional activity of MOR was assessed by the [^35^S]-GTPγS binding assay, as described previously [21]. The method is based on the ability of the radiolabeled GTP analog, [^35^S]-GTPγS, to specifically bind to G-proteins. Here, 100mg of AI tissue was used for each subject for membrane preparation. Membranes were prepared by homogenizing the tissue in ice-cold 0.25 M sucrose solution 10 vol (ml/g wet weight of tissue). Samples were then centrifuged at 1100 g for 10 min. Supernatants were collected and diluted 5 times in buffer containing 50 mM Tris-HCl, pH 7.4, and 1 mM EDTA, and were then centrifuged at 35,000 g for 30 min. The pellets were homogenized in 2 ml of ice-cold sucrose solution (0.32 M), aliquoted and kept at −80°C until further use. For each [^35^S]-GTPγS binding assay, 2.5 μg of membrane proteins were used per well. Samples were incubated with the MOR agonist DAMGO (using 12 increasing concentrations, from 10^−4^ to 10^−10^ M) for 1 h at 25 °C in assay buffer 50 mM Tris–HCl (pH 7.4), 3 mM MgCl_2_, 100 mM NaCl, 0.2 mM EGTA containing 30 μM GDP and 0.1 nM [^35^S]-GTPγS. Incubation was terminated by rapid filtration and washing in ice-cold buffer (50 mM Tris–HCl, 5 mM MgCl_2_, 50 mM NaCl, pH 7.4). Bound radioactivity was quantified using a liquid scintillation counter (Perkin Elmer Top Count). Non-specific binding was defined as binding in the presence of 10 μM GTPγS, and basal binding was assessed in the absence of DAMGO. Assays were performed in triplicates in different membrane preparations. Non-linear regressions were performed on individual concentration-response curves of DAMGO-induced stimulations (Prism, GraphPad Software, San Diego, CA, USA). Data are expressed as the maximal net stimulation (E_max_, stimulated minus basal levels) expressed in fmol.mg^−1^ protein. The concentration that elicited half-maximal effect (EC_50_) was obtained and normalized as log EC_50_ values.

### Quantification of inhibitory G-proteins alpha subunits expression using Nanostring

Nanostring experiments were performed as described previously [22] at the Jewish General Hospital Molecular Pathology Centre (Montréal, QC, Canada) using Nanostring nCounter targeted gene expression profiling [23]. Briefly, 5 μl of total RNA (20 ng/μl) was hybridized with the reporter and capture probes at 65^°^C in a thermocycler for 19-20 hours. Probes were designed against 4 genes of interest (GNAI1, GNAI2, GNAO1 and GNAZ) and against 4 housekeeping control genes highly expressed in the AI: GAPDH, β-Actin, Tubulin and β-2-Globulin. The samples were processed with the nCounter Prep Station to purify the hybridized targets and affix them to the cartridge. After transfer to the nCounter Digital Analyzer, barcodes were counted and tabulated for each target molecule. The data were analysed using the nSolver version 2.6.43 following the manufacturer’s recommendations.

### Statistical Analysis

Statistical analyses were carried out using R and Prism v8.4.1. Frequencies of SNPs were analysed using *χ*^*2*^ test to test for Hardy-Weinberg equilibrium. For MOR expression and [^35^S]-GTPγS a general linear model (GLM) was used to analyse group differences. Potential correlations between variables that commonly act as confounders (pH, PMI, RIN, and age) and gene expression were assessed separately, and only those variables showing a significant Pearson correlation were included in each final GLM model. Statistical significance was set at p<0.05.

## Results

### MOR gene expression in the anterior insula

We first used qPCR to quantify MOR mRNA levels in the two groups of C and MDD subjects. Groups did not differ with respect to the subjects’ sex (Fisher’s exact test, p>0.99) nor RNA Integrity Numbers (RIN; t=1.25, p=0.21; see Table1). MOR expression was similar across men and women (t=1.30, p=0.20) and, as expected, significantly correlated with RIN values (Fig.S1A). Interestingly, it also increased with age (Fig.1A; r=0.37, N=90 p=3E-04), in contrast with previous studies that reported a significant decrease in the medio-dorsal thalamus [19], with no changes in three other structures (the caudal part of the anterior cingulate cortex, cACC [19], the ventral posterior part of the lateral thalamus [24], and the secondary somato-sensory cortex [24]). This could indicate brain region-specific patterns of MOR regulation during aging. Importantly, a GLM comparing our two main groups, C and MDD, and considering age and RIN as significant covariates, did not reveal a difference between groups (Fig.1B, F(1,86)=1.11, p=0.27). Further, while we previously reported that a history of child abuse, a major distal risk factor for MDD, associated in the AI with decreased expression of the kappa opioid receptor [19], no such effect was found for MOR (t=0.83, p=0.41).

**Figure 1.**
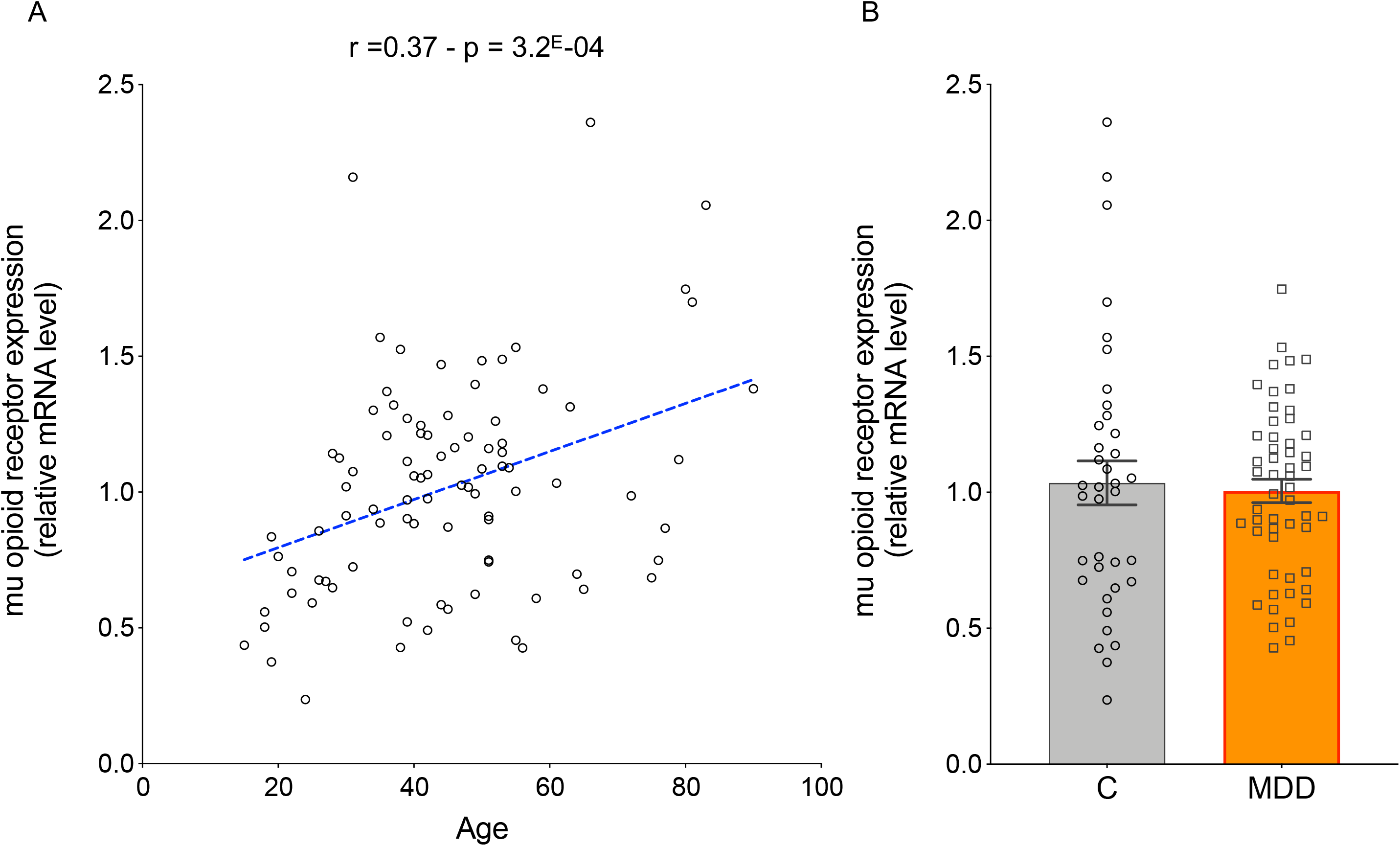
Relative real-time quantitative PCR (RT-PCR) expression of the mu opioid receptor (MOR) in the anterior insula by age and clinical group. **A**. MOR expression was quantified by RT-PCR in quintuplicates, using calibration curves and two reference housekeeping genes: GAPDH and β-actin (see methods for details). The scatter plot displays a significant positive linear correlation between MOR expression and age, in all subjects, independent of clinical group (r=0.37, p=3E-04). **B**. Expression of MOR in healthy controls and subjects who died during a major depressive episode (MDD). There was no significant difference between the two clinical groups (F(1,86)=1.11, p=0.27). Mean ± sem, as well as individual expression values, are shown.

All drugs of abuse, including opiates, trigger rewarding effects at least in part by recruiting the MOR [25], which has been conceptualized as a molecular gateway to addiction [26]. Animal studies have also reported changes in MOR expression following acute or chronic activation of this receptor. We therefore explored whether MOR expression varied as a function of comorbid substance use disorder (SUD), or ante-mortem acute opiate consumption. Results showed that the SUD diagnosis, present at similar frequency in C and MDD groups (Fisher’s exact test, p>0.99; Table 1), had no significant effect (Fig.S1B, t=0.32, p=0.75). Toxicological screenings (conducted on blood, urine, and ocular liquid samples) indicated that all drugs of abuse considered together (opiates, cocaine, amphetamines, ethanol, and tetrahydrocannabinol), or opiates only, were detected in 35 and 8 subjects, respectively, and significantly associated with SUD diagnosis, as expected (p<0.05). However, positive toxicological screening did not associate with any change in MOR expression (p>0.05; Fig.S1C). Altogether, MOR expression in the AI was not modulated by MDD, SUD, recent opiate consumption, or history of child abuse.

### MOR gene expression and single nucleotide polymorphisms (SNP)

We investigated a series of SNP in the MOR gene (see observed frequencies in Table 2). These were chosen based on previous literature [24,27,28], and notably included the A118G (rs1799971), a well-studied SNP in relation to MOR function and SUD [29]. The distribution of the A and G alleles for A118G was in Hardy-Weinberg equilibrium (HWE), as assessed using a *χ*^*2*^ test (*χ*^*2*^=0.52, p=0.47), with a low frequency of the G allele (approx. 13.6%), as expected in a Quebec population of European descent, and similar to previous studies [30,31]. When pooling results from the present work with those previously obtained in two other brain regions (the caudal part of the anterior cingulate cortex and medio-dorsal thalamus [19]), the A118G genotype was found to have a moderately significant effect on MOR mRNA expression, with higher expression in carriers of the minor G allele (Fig.S2; 2-way ANOVA: genotype effect F(1,184)=6.30, p=0.013; brain region effect F(2,184)=1.53, p=0.22). Previous studies that investigated this same SNP reported no correlation with MOR expression, neither in the thalamus nor the somato-sensory [24] or dorsolateral prefrontal cortices [32], suggesting a brain region-specific impact of this SNP. For the 15 other SNPs that we investigated, no relationship with MOR expression was observed (p>0.05 for all).

**Table 2.** Single nucleotide polymorphisms (SNP) profiled in this study in the *OPRM1* gene encoding the mu opioid receptor. SNPs were determined using Sanger sequencing (see methods). Genomic position (in the Human GRCh37/hg19 reference genome), gene feature (promoter/exon/intron), observed counts and percentages of major and minor alleles in our cohort, and expected percentages (according to the *1000 Genomes Project*) are shown.

| SNP | Position | Feature | Sample Count<br>Major / Minor | Sample Frequency<br>Major / Minor (%) | 1000 Genomes frequency<br>Major / Minor (%) |
| --- | --- | --- | --- | --- | --- |
| rs147391824 | chr6:154,348,457 | Promoter | A (179) / C (7) | A (96.237) / C (3.763) | A (98.482) / C (1.518) |
| rs7751510 | chr6:154,348,488 | Promoter | G (176) / C (10) | G (94.624) / C (5.376) | G (89.317) / C (10.683) |
| rs117514803 | chr6:154,348,559 | Promoter | C (172) / T (14) | C (92.473) / T (7.527) | C (98.003) / T (1.997) |
| rs7776341 | chr6:154,348,605 | Promoter | A (177) / C (9) | A (95.161) / C (4.839) | A (89.357) / C (10.643) |
| rs1074287 | chr6:154,348,809 | Promoter | A (137) / G (49) | A (73.656) / G (26.344) | A (71.825) / G (28.175) |
| rs60509363 | chr6:154,348,978 | Promoter | G (177) / A (9) | G (95.161) / A (4.839) | G (85.903) / A (14.097) |
| rs17181003 | chr6:154,360,444 | Exon 1 | A (184) / G (0) | A (100) / G (0) | A (98.942) / G (1.058) |
| rs6912029 | chr6:154,360,508 | Exon 1 | G (175) / T (9) | G (95.109) / T (4.891) | G (92.712) / T (7.288) |
| rs17174638 | chr6:154,360,569 | Exon 1 | C (184) / T (0) | C (100) / T (0) | C (96.745) / T (3.255) |
| rs1799972 | chr6:154,360,696 | Exon 1 | C (184) / T (0) | C (100) / T (0) | C (92.871) / T (7.129) |
| rs1799971 | chr6:154,360,797 | Exon 1 | A (159) / G (25) | A (86.413) / G (13.587) | A (77.656) / G (22.344) |
| rs12209447 | chr6:154,361,048 | Intron 1 | C (175) / T (9) | C (95.109) / T (4.891) | C (96.046) / T (3.954) |
| rs9479757 | chr6:154,411,344 | Intron 2 | G (164) / A (16) | G (91.111) / A (8.889) | G (91.953) / A (8.047) |
| rs17181199 | chr6:154,411,710 | Intron 2 | T (184) / A (0) | T (100) / A (0) | T (98.642) / A (1.358) |
| rs17181213 | chr6:154,411,847 | Intron 2 | C (186) / T (0) | C (100) / T (0) | C (98.502) / T (1.498) |
| rs2075572 | chr6:154,412,004 | Intron 2 | C (113) / G (67) | C (62.778) / G (37.222) | C (61.162) / G (38.838) |

### MOR signaling efficiency

We next used [^35^S]-GTPγS to determine MOR G-protein coupling. Using DAMGO as a well-validated agonist, we generated dose-response curves for each subject, which were then used to compute maximal receptor stimulation (E_max_) and the agonist concentration eliciting half-maximal effect (log EC_50_).

First, we found a significant correlation between inter-individual variability of MOR mRNA (qPCR) and protein (MOR E_max_) levels (r=0.28, p=0.028; Fig.2A). This suggests that signaling of the receptor in the AI is at least partly explained by its expression from local neurons, and provides a validation of our [^35^S]-GTPγS procedure. Second, we found that basal [^35^S]-GTPγS binding (defined as the constitutive activity of all GPCRs, measured in the absence of DAMGO), was slowly decreasing as a function of age (r=(−0.23), p=0.028; Fig.S3A), consistent with previous reports [33]. MOR E_max_ was not affected by either sex (p=0.42) or age (p=0.31), but significantly correlated with brain tissue pH (r=0.34, p=7.2E-05; Fig.S3B). Finally, we observed that MOR affinity for DAMGO, measured as the log EC_50_, showed a significant negative correlation with age (r=−0.31, p=0.0023; Fig.2B), corresponding to a leftward shift of concentration-response curves, and indicating enhanced DAMGO efficacy with aging. This is consistent with results from a previous study conducted in the prefrontal cortex (Broadmann Area 9, see [33]), although the biological significance of this effect is elusive.

**Figure 2.**
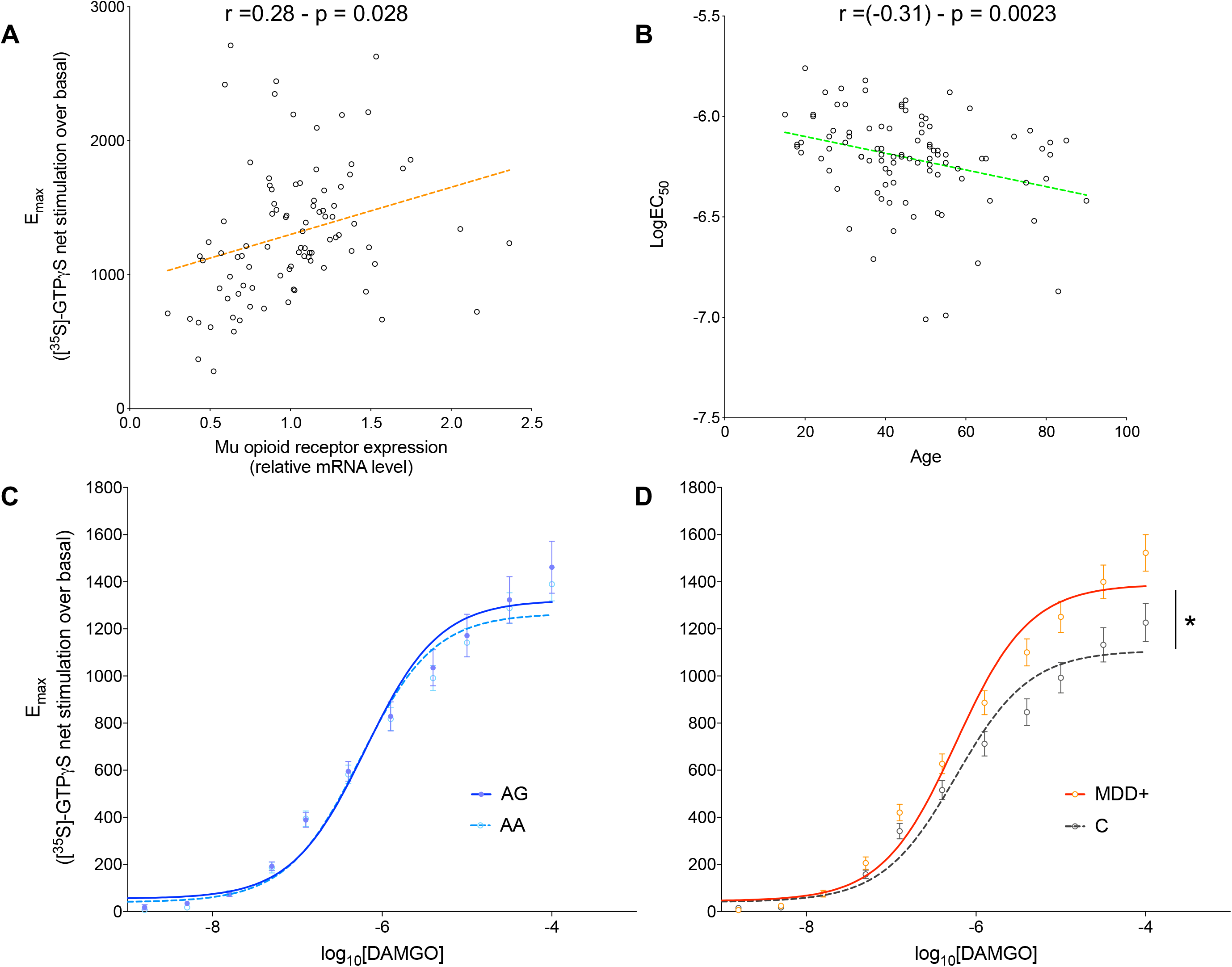
Mu opioid receptor (MOR) DAMGO agonist-stimulated [^35^S]-GTPγS binding in the anterior insular cortex of healthy controls and subjects with major depressive disorder (MDD). Coupling efficiency of MOR was evaluated using [^35^S]-GTPγS binding and 12 increasing concentrations of the potent and specific agonist DAMGO, followed by non-linear regression of concentration-response curves, to compute E_max_ (maximal net stimulation of MOR; [^35^S]-GTPγS net stimulation over basal binding) and logEC_50_ (agonist concentration for half-maximal effect, log transformed) values. **A**. The scatter plot displays a significant positive linear correlation (r=0.28, p=0.028) between MOR mRNA levels (measured by RT-PCR) and protein signaling (E_max_). **B**. Scatter plot displaying a significant negative linear correlation (r=(−0.31), p=0.0023) between age and MOR affinity for DAMGO (logEC_50_). **(C-D)** Concentration-response curves of [^35^S]-GTPγS stimulation, by log_10_[DAMGO], in the anterior insular cortex. **C**. Dose-response curve in non-carriers (broken line; AA genotype) or carriers of the minor G allele (filled-in line; AG genotype) for the MOR single-nucleotide polymorphism rs1799971 (A118G). The two genotypes did not show any significant differences in MOR E_max_ (p=0.64) or logEC_50_ (p=0.97) values. **D**. Dose-response curve in control (broken line) or MDD (filled in line) subjects. Subjects diagnosed with MDD showed a significant increase in MOR coupling efficiency in the anterior insular cortex (F(1,94)=4.97, p=0.028; controlling for the significant effect of pH, see main text). Values are mean ± sem. * p <0.05.

We next wondered whether MOR coupling was modulated by genetic variation, as per our SNP data. We used dominant negative models and linear regression to investigate relationships between MOR E_max_ and observed allelic frequencies. Results showed a nominally significant correlation for rs1074287, a SNP located upstream the canonical MOR transcription start site and within the first intron of rare transcript variants (see Fig.S4A). Accordingly, a higher potency of the DAMGO agonist was observed in carriers of the minor G allele (p=0.017; Fig.S4B), suggesting higher MOR protein abundance in these subjects. This effect, however, did not stand multiple testing correction (n=16 SNPs; Bonferroni-corrected p-value=0.27), and will require replication in a larger cohort. In addition, no significant effect of the minor G allele was observed on MOR expression (p=0.88; Fig.S4C). For other SNPs investigated, no significant correlation was observed (p>0.05; data not shown), including for the A118G SNP (Fig.2C). Regarding the later, to our knowledge, two previous studies similarly compared MOR coupling across A and G carriers [24,34], both using brain tissue dissected from the secondary somatosensory cortex (SII) and ventral posterior part of the lateral thalamus. The first study [34] (total sample size N=33) found lower DAMGO efficacy in G carriers in the SII, but not in the thalamus, while the second study [24] (total sample size N=60), conducted in another cohort, found lower efficacy in G carriers in both brain regions. Combined with our findings, this suggests that the A118G might exert a brain-region specific effect on MOR coupling. This hypothesis is further supported by results from an animal study conducted in mice genetically modified to model the human polymorphism, which reported similar brain-region specificity (with no significant effect of genotype in the AI, see [35]).

Finally, we investigated MOR coupling in relation to psychopathology. Similar to aforementioned analysis of MOR mRNA expression, we first wondered whether the [^35^S]-GTPγS assay was impacted by ante-mortem consumption of drugs of abuse, or SUD diagnosis. Results showed that none of these factors had any effect on MOR E_max_ (SUD: p=0.55, FigS3C; acute opiate: p=0.74, FigS3D). We then built a GLM model to search for an association with MDD, taking into account the effect of pH as covariate (see above and FigS3B). Results indicated that depressed subjects showed significantly higher E_max_ (F(1,94)=4.97, p=0.028; Fig 2D, an effect that remained significant even when correcting for the effect of rs1074287; p=0.0447). This was not accompanied by any change in logEC_50_ as a function of MDD (p=0.58), indicating that MOR protein signaling, but not affinity, is affected by MDD status. Overall, these results show that coupling of the MOR to inhibitory G-proteins is potentiated in individuals diagnosed of MDD, independently of comorbid SUD.

### Expression of inhibitory alpha G-protein subunits

MOR belongs to the family of inhibitory G-protein coupled receptors that activate Gi alpha-subunits, thereby inhibiting adenylate cyclase activity and lowering cAMP levels. To determine whether changes in expression of these proteins might contribute to altered MOR coupling in MDD subjects, we used Nanostring to quantify the expression of the four major corresponding genes: GNAI1, GNAI2, GNAO1, and GNAZ. We first looked at potential effects of clinical and technical covariates, and found none for sex or SUD diagnosis (p>0.05), while RIN affected GNAI1 only (p=0.029). Of note, aging associated with a significant decrease in GNAZ expression (p=0.016; FigS5), an effect that may contribute to the decrease in basal binding (i.e. in the absence of MOR agonist) previously observed in [^35^S]-GTPγS experiments (see above, and FigS4A). More importantly, we investigated potential association with MDD, and observed a selective increase in the expression of GNAI2 in depressed individuals (GLM, F(1,89)=7.42, p=0.0077; Fig3), which remained significant even when controlling for the number of genes investigated by Nanostring (Bonferroni-corrected p-value=0.031). Therefore, our results suggest that increased expression of the GNAI2 alpha subunit is a molecular adaptation that may contribute to enhanced MOR coupling in subjects with MDD.

**Figure 3.**
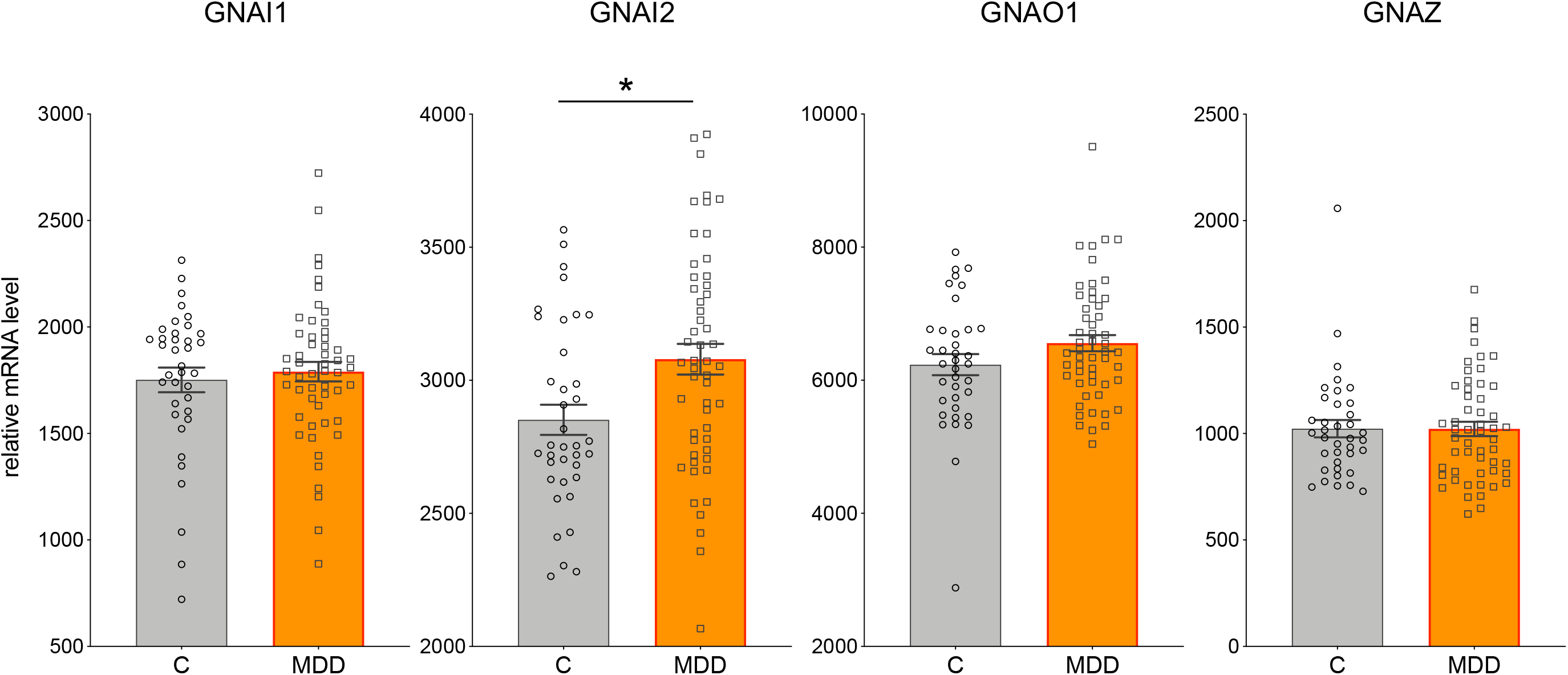
Relative expression of major inhibitory alpha G-protein subunits in the anterior insula of healthy controls and subjects with major depressive disorder (MDD). NanoString was used to quantify the mRNA relative expression of 4 inhibitory alpha G-protein subunits. No difference was found between controls and depressed individuals for 3 subunits: GNAI1, GNAO1 and GNAZ. In contrast, subjects with MDD showed a significant increase in the expression of the GNAI2 subunit, compared to controls (F(1,89)=7.42, p=0.0077). Mean +/− SE, as well as, individual expression values are shown, * p <0.05.

## Discussion

Inspired by recent imaging studies reporting convergent evidence for MOR dysfunction in the AI of individuals with MDD, the present post-mortem study investigated its molecular expression and function at this brain site.

We first quantified MOR gene expression using qPCR, and found no significant change as a function of MDD. To our knowledge, only two previous studies quantified MOR mRNA in depressed individuals compared to healthy controls, and reported increased levels in the prefrontal cortex (BA9, see [36]), or no change in the mediodorsal thalamus or anterior cingulate cortex (BA24/32, [19]), altogether suggesting that MDD-related changes in MOR expression may be subtle, and specific to particular brain regions.

We then used [^35^S]-GTPγS to assess MOR signalling. In the past, four human studies have investigated MOR density using the radiolabelled agonist [^3^H]-DAMGO [9,11–13], while only one study used agonist-stimulated [^35^S]-GTPγS to measure both receptor affinity and density [21]. The two oldest studies, although conducted in small cohorts ([11], N=2-7 subjects/group; [12], N=15/group), provided preliminary evidence for increased MOR abundance in the frontal and temporal cortices, and the caudate nucleus. However, the three subsequent studies, performed on frontal cortex tissue (BA9 and BA24) in larger cohorts ([9], N=9-10/group; [21], N=28/group; [13], N=20-51/group), failed to replicate these findings, as they all reported no significant differences between subjects who died by suicide (mostly diagnosed with mood disorders) and controls. In the present study, we investigated a larger cohort (N=40-55/group) and focused instead on the AI, a brain region previously not investigated using such assays. Our findings indicate that MDD was associated in the AI with a significant increase in MOR activity, an effect that occurred in the absence of any change in receptor affinity, and that was not accounted for by other confounders such as opiate consumption, SUD, or clinical covariates. This suggests, therefore, that the AI is a prominent brain site where MDD-related changes affecting MOR may contribute to disease pathophysiology.

To understand molecular adaptations potentially underlying enhanced MOR coupling, and because MOR is an inhibitory G-protein-coupled receptor, we next used Nanostring to quantify expression of the 4 major inhibitory alpha G-protein subunits. Results showed a specific increase in the expression of GNAI2 in depressed individuals. Because [^35^S]-GTPγS Emax measures at saturating agonist concentration reflect maximal signalling when all receptors are recruited, this increased GNAI2 expression likely explains the increase in Emax detected in the GTPgS assay.

Importantly, these findings appear consistent with imaging studies published over the last 15 years (see [37] for review), mostly conducted using positron emission tomography (PET) and the MOR selective radiotracer [^11^C]-Carfentanil. These studies first indicated that acute emotionally salient stimuli (whether the recall of sad autobiographical events [4], exposure to social rejection [7], or viewing pictures of appetizing food [38]) led in heathy volunteers and depressed individuals [8] to rapid changes in MOR availability, notably at the level of the AI, reflecting the well-characterized notion that MOR activity modulates hedonic tone.

Beyond these fluctuations, prolonged low mood in MDD also seems to affect MOR function. Accordingly, while an initial PET study reported lower MOR availability in depressed women, which reached significance in the posterior thalamus only (possibly due to modest statistical power, N=14/group) [5], a more recent study found convergent results by investigating a large cohort characterized for subclinical depressive symptoms (N=135, with 113 males; see [2]). Widespread reductions in MOR availability were observed as a function of symptom severity across numerous cortical regions, with the insula, precuneus, and temporoparietal regions most strongly affected. Importantly, such reduced availability can be interpreted in PET experiments as reflecting either increased occupancy of the MOR by endogenous opioid peptides or, alternatively, lower receptor density or affinity. Results from the present post-mortem study argue in favor of the former possibility. Because increased MOR signaling was found in our study, we hypothesize that depression associates in the AI with higher MOR coupling efficiency (Emax) that at least partly stem from higher GNAI2 expression. In addition, PET studies suggest enhanced endogenous release of opioid peptides (lower MOR availability). Therefore, these 2 distinct mechanisms seem to converge to potentiate the insular opioidergic tone, and likely reflect a compensatory mechanism that counteracts low mood.

While MOR-mediated regulation of reward and mood has been primarily investigated in animal models in brain structures associated with the mesolimbic dopaminergic system, such as the ventral tegmental area and basal ganglia [1,39], it has been left relatively unexplored in cortical structures. Studies in human, however, clearly indicate that MOR signalling in the AI contributes to important physiological function, in particular the processing of emotional aspects of pain that may arise from either somatosensorial stimuli (physical pain), or from situations that threatens the organism’s survival, internal homeostasis, or social position (i.e. psychological or social pain) [40,41]. Our findings are consistent with this hypothesis, and suggest that opiate medications that recruit the MOR (e.g. buprenorphine, frequently combined with samidorphan), and are currently under intense examination for the management of depression and suicidal ideas [42,43], may act at least in part by potentiating in the AI the beneficial impact of MOR on social and emotional distress.

In conclusion, the present work brings new knowledge that, in combination with imaging studies, indicates that the AI is a critical brain site where increased MOR signalling occurs in the context of MDD and suicide. Future studies will be necessary to determine whether similar adaptations may affect other brain regions where MOR contributes to reward and affective regulation.

## Contributions

PEL, FJ, BLK and GT conceived the study. PEL performed RNA extractions, qPCR, and Nanostring experiments. DA conducted SNP genotyping. DF, PEL and BLK conceived and performed [^35^S]-GTPγS experiments. PEL prepared the manuscript, and all authors approved its final version.

