## Supplementary Figures for "Increased functional coupling of the mu opioid receptor in the anterior insula of depressed individuals"

Gustavo Turecki, MD PhD

McGill Group for Suicide Studies, Douglas Mental Health Institute

Department of Psychiatry, McGill University

#### Content

. Legends of supplementary figures

. Supplementary Figures

#### Supplementary Figures

**Figure S1. Relative qPCR expression of the mu opioid receptor (MOR) in the anterior insula. A.** Scatter plot displaying a significant positive correlation between MOR expression and RIN, in all subjects independent of group (r =0.36, p = 5.4E-04). **B.** Expression of MOR was similar (p=0.75) in subjects with or without a diagnosis of substance use disorder (SUD). **C.** Similarly, no significant change in MOR expression (p>0.05) was found between subjects with a negative (circles) or positive (squares) toxicological screening result for all drugs of abuse (left), or opiates alone (right). Mean +/- SE, as well as, individual expression values are shown.

**Figure S2. Relative qPCR expression of the mu opioid receptor (MOR) across brain regions by rs1799971 genotype.** Subjects were separated into non-carriers (circles) or carriers of the minor G allele (squares) of the rs1799971 single nucleotide polymorphism (A118G), and MOR mRNA expression levels quantified by qPCR in the anterior insula (left), caudal anterior cingulate cortex (middle), and medio-dorsal thalamus (right). Carriers of the minor G allele showed higher expression, with no differences among brain regions, and no interaction between factors. (2-way ANOVA: genotype effect, $F(1,184)=6.30$, p=0.013; brain region effect, $F(2,184)=1.53$, p=0.22). Mean +/- SE, as well as, individual expression values are shown.

**Figure S3. [$^{35}$S]-GTPγS stimulation of the mu opioid receptor (MOR), using the DAMGO agonist, in the anterior insular cortex. A.** Scatter plot displaying a significant negative correlation (r=(-0.23), p=0.028) between age and basal [$^{35}$S]-GTPγS binding (defined as the constitutive activity of all GPCRs, measured in the absence of DAMGO). **B.** Scatter plot displaying a significant positive correlation (r=0.34, p=7.2E-05) between pH and MOR levels ([$^{35}$S]-GTPγS net over basal stimulation). **C.** MOR signaling efficiency ([$^{35}$S]-GTPγS net over basal stimulation) was similar (p=0.55) across subjects with or without a diagnosis of substance use disorder (SUD) **D.** Similarly, no difference (p=0.74) was observed in MOR signaling ([$^{35}$S]-GTPγS net over basal stimulation) between subjects with a negative (circles) or positive (squares) toxicological screening result for all drugs of abuse (left), or opiates alone (right).

**Figure S4. Effects of rs1074287 genotype on mu opioid receptor (MOR) mRNA levels and signaling in the anterior insular cortex. A.** Location of rs1074287, a SNP located upstream the canonical MOR transcription start site and within the first intron of rare transcript variants. **B.** Dose-response curve of [$^{35}$S]-GTPγS stimulation, by $\log_{10}$[DAMGO], showing increased MOR signaling in carriers of the minor G allele (filled in line) compared to non-carriers (broken line) of the rs1074287 polymorphism (p=0.017). **C.** This polymorphism, however, did not associate with any change in MOR mRNA expression, which was similar across non-carriers (broken line) or carriers of the minor G allele (filled in line; p=0.88). * $p < 0.05$

**Figure S5. Expression of *GNAZ* in the anterior insular cortex.** The scatter plot displays a significant negative correlation between relative NanoString expression of *GNAZ* in the anterior insula and age, in all subjects, independent of group (r=(-0.25), p=0.016).

**Figure S1**

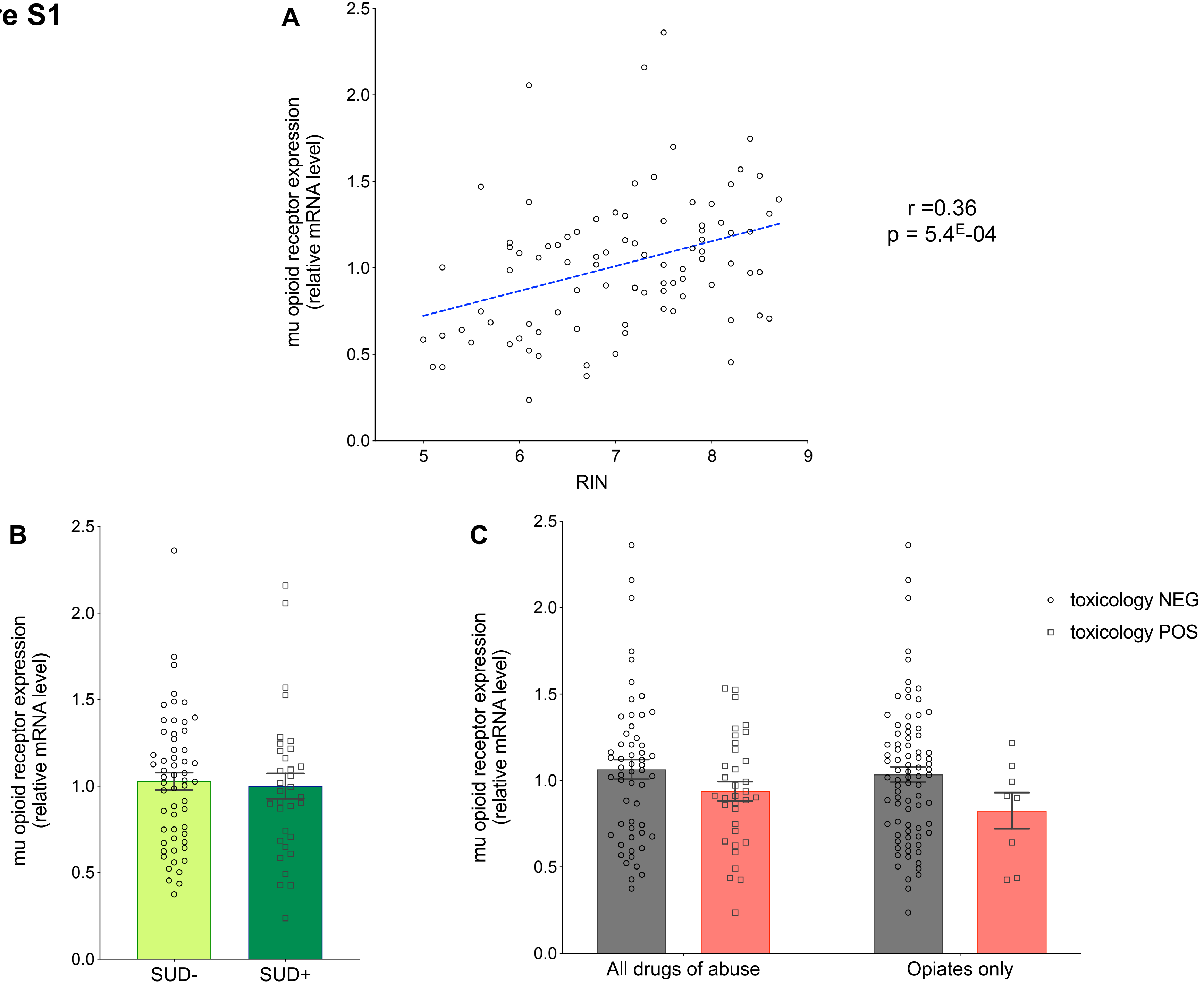

A
mu opioid receptor expression
(relative mRNA level)
2.5
2.0
1.5
1.0
0.5
0.0
5
6
7
8
9
RIN
r =0.36
p = 5.4E-04
B
mu opioid receptor expression
(relative mRNA level)
SUD-
SUD+
C
mu opioid receptor expression
(relative mRNA level)
All drugs of abuse
Opiates only
toxicology NEG
toxicology POS

**Figure S2**

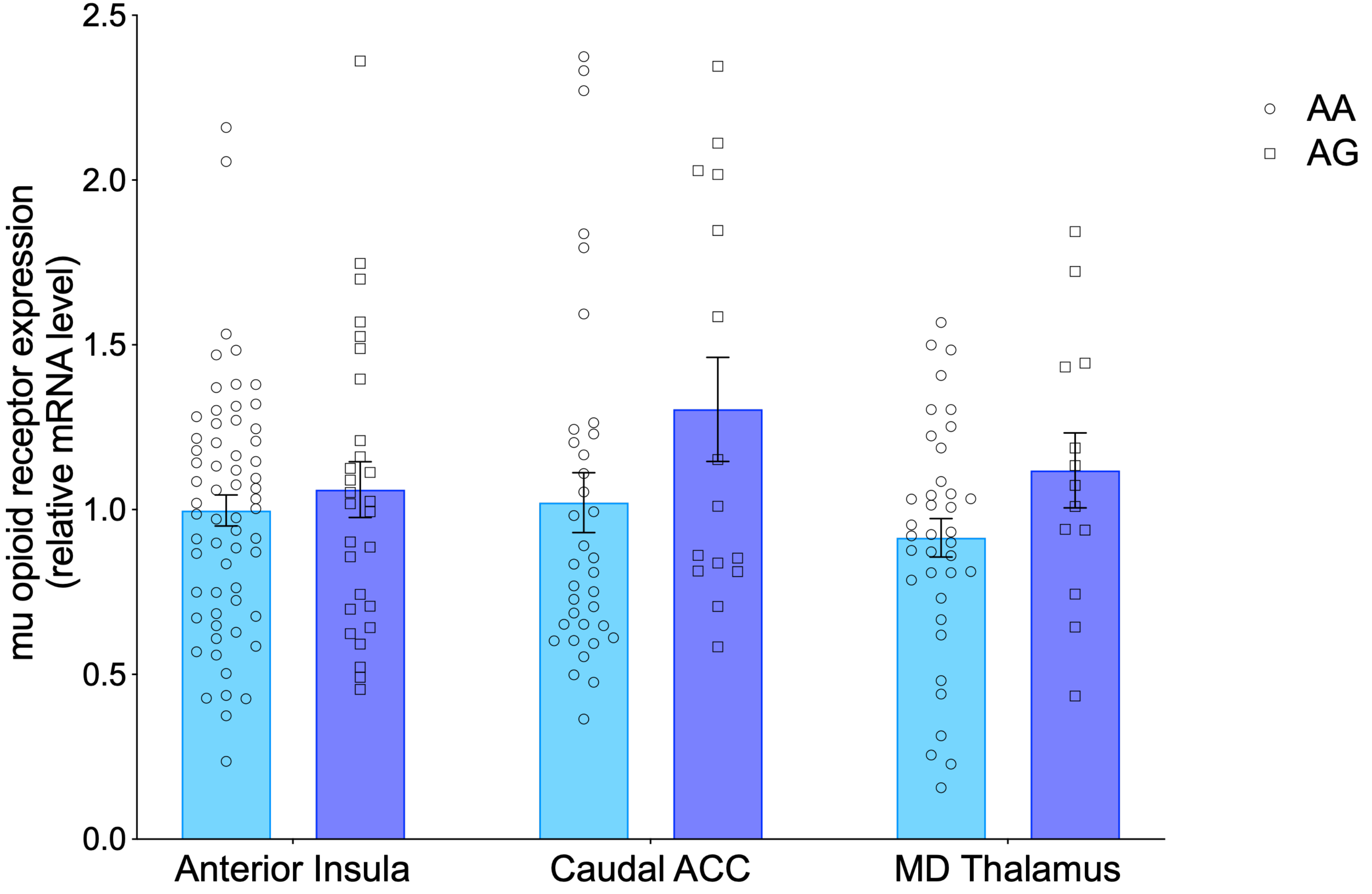
mu opioid receptor expression
(relative mRNA level)
2.5
2.0
1.5
1.0
0.5
0.0
Anterior Insula
Caudal ACC
MD Thalamus
AA
AG

### Figure S3

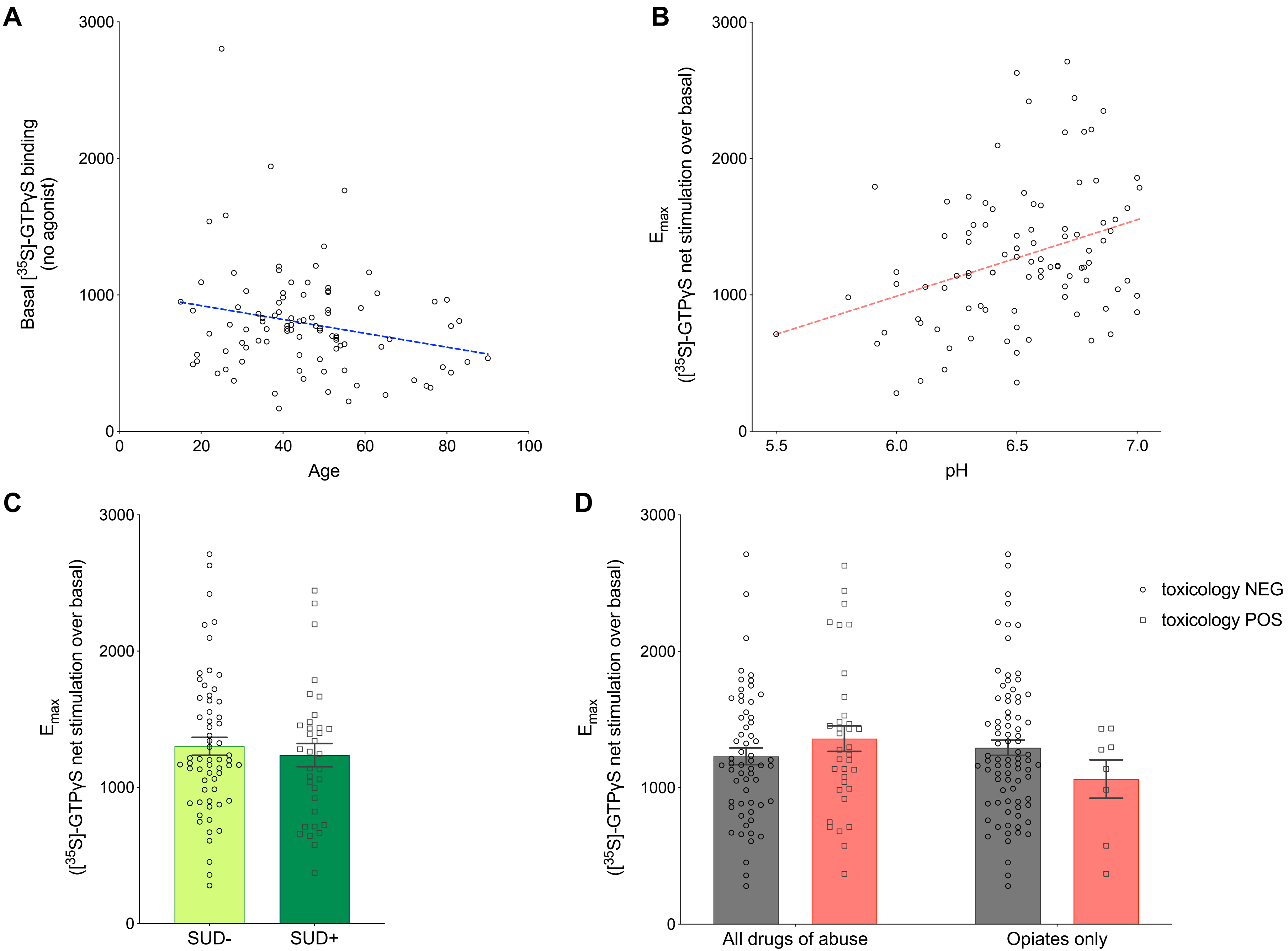

A
Basal [35S]-GTPγS binding (no agonist)
3000
2000
1000
0
0
20
40
60
80
100
Age
B
Emax ([35S]-GTPγS net stimulation over basal)
3000
2000
1000
0
5.5
6.0
6.5
7.0
pH
C
Emax ([35S]-GTPγS net stimulation over basal)
3000
2000
1000
0
SUD-
SUD+
D
Emax ([35S]-GTPγS net stimulation over basal)
3000
2000
1000
0
All drugs of abuse
Opiates only
toxicology NEG
toxicology POS

Figure S4

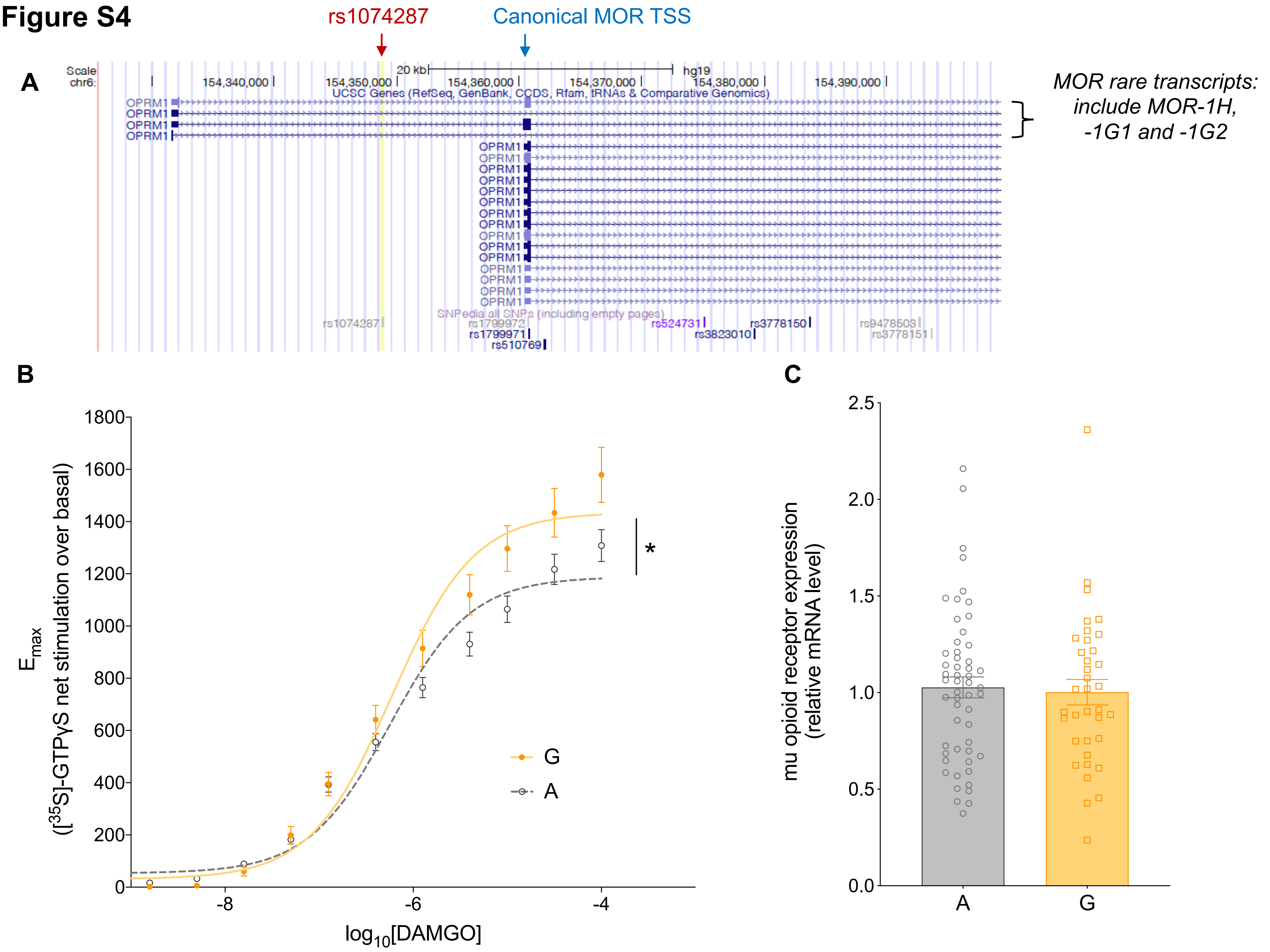
A
rs1074287
Canonical MOR TSS
Scale
chr6:
20 kb
hg19
154,340,000
154,350,000
154,360,000
154,370,000
154,380,000
154,390,000
UCSC Genes (RefSeq, GenBank, CCDS, Rfam, tRNAs & Comparative Genomics)
OPRM1
SNPedia all SNPs (including empty pages)
rs1074287
rs1799972
rs1799971
rs510769
rs524731
rs3823010
rs3778150
rs9478503
rs3778151
MOR rare transcripts: include MOR-1H, -1G1 and -1G2
B
Emax
([35S]-GTPγS net stimulation over basal)
log10[DAMGO]
G
A
*
C
mu opioid receptor expression (relative mRNA level)
A
G

**Figure S5**

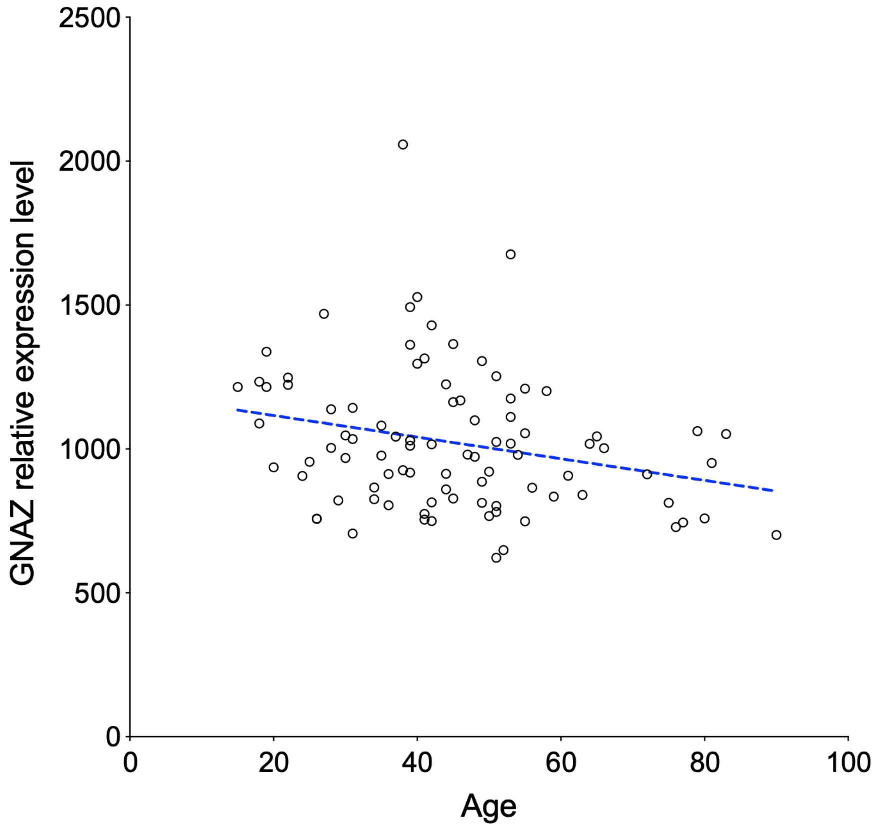

2500
2000
1500
1000
500
0
GNAZ relative expression level
0
20
40
60
80
100
Age
